# High-throughput cellular-resolution synaptic connectivity mapping *in vivo* with concurrent two-photon optogenetics and volumetric Ca^2+^ imaging

**DOI:** 10.1101/2020.02.21.959650

**Authors:** Christopher McRaven, Dimitrii Tanese, Lixia Zhang, Chao-Tsung Yang, Misha B. Ahrens, Valentina Emiliani, Minoru Koyama

**Affiliations:** Janelia Research Campus, HHMI, Ashburn VA 20147, USA; Institut de la Vision, Sorbonne University, Inserm S968, CNRS UMR7210, 17 Rue Moreau, 75012, Paris, France

## Abstract

The ability to measure synaptic connectivity and properties is essential for understanding neuronal circuits. However, existing methods that allow such measurements at cellular resolution are laborious and technically demanding. Here, we describe a system that allows such measurements in a high-throughput way by combining two-photon optogenetics and volumetric Ca^2+^ imaging with whole-cell recording. We reveal a circuit motif for generating fast undulatory locomotion in zebrafish.

## Main

Synapses conduct electrical signals between neurons and allow them to form circuits that ultimately control animal behavior. Thus, the ability to map synaptic connectivity and synaptic properties is essential for understanding circuit operations underlying animal behavior. Electrical signals through synapses and the dynamics of these signals can be directly measured with electrophysiological techniques such as paired whole-cell recordings. However, these techniques are hard to scale up and the yield is often low, especially *in vivo*. The development of optogenetics enabled optical control of neuronal activity has been used to assess macroscopic synaptic connectivity wherein a group of neurons is activated and their synaptic responses in other neurons are monitored via whole-cell recordings^1^. The next step is to transition from this macroscopic approach into a microscopic synaptic connectivity mapping. This requires performing high throughput optogenetic activation of presynaptic cells with single cell resolution while monitoring the photo-evoked activity of the presynaptic cell. The combination of two-photon shaped stimulation and the engineering of efficient soma-targeted opsins, called circuits optogenetics^2^, opened a way to stimulate single and multiple neurons at near single-cell resolution^3–5^. Yet, the effectiveness of this approach for synaptic connectivity mapping has only been demonstrated in proof-of-principle experiments involving relatively small neuronal populations in brain slices^3,5^ and has not been exploited for large-scale synaptic connectivity mapping *in vivo*. Furthermore, potential 3D off-target activations and cross talk excitation of the opsins from the imaging laser have never been rigorously assessed for such connectivity analyses.

Here we present a concurrent two-photon optogenetic stimulation and volumetric Ca^2+^ imaging system for zebrafish that allows one to induce and monitor single-cell activation in 3D and reconstruct cellular-resolution synaptic property maps in a large volume *in vivo* (Fig. 1ai), which is prohibitively difficult to accomplish with traditional techniques. We examine long-range synaptic connectivity of genetically defined spinal interneurons and reveal a circuit motif that may underlie fast undulatory movements during rapid forward locomotion. To our knowledge, this is the first demonstration of high-throughput physiological measurements of synaptic properties at single-cell resolution *in vivo* that provides insights into circuit operations in the nervous system.

**Figure 1.**
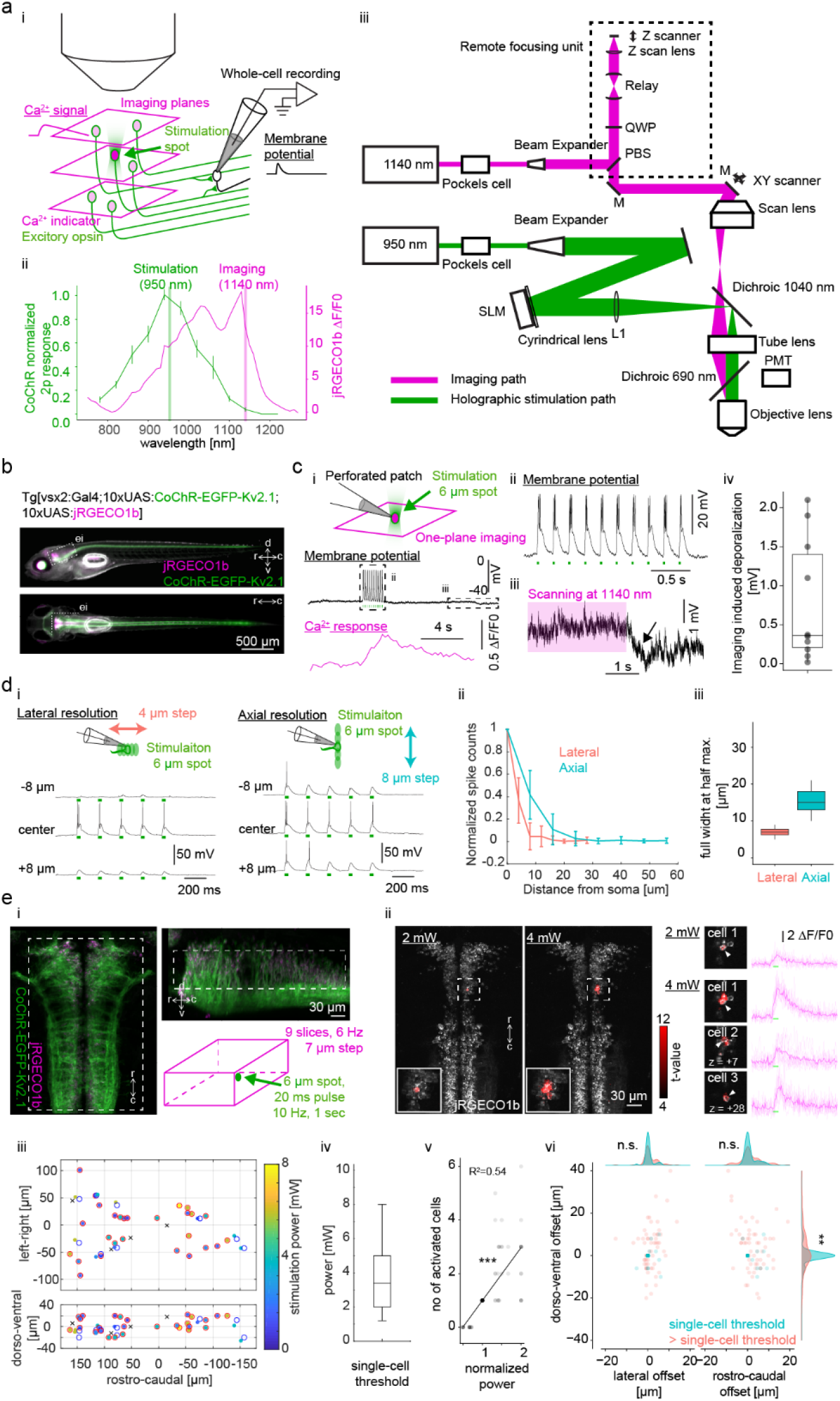
Concurrent two-photon optogenetics and volumetric Ca^2+^ imaging for high-throughput synaptic connectivity mapping. **a.** (i) A schematic of the high-throughput cellular synaptic connectivity mapping using two-photon optogenetics and volumetric imaging. A genetically defined cell group is co-labeled with Ca^2+^ indicator (magenta) and excitatory opsin (green). Multiplane imaging is used to monitor the 3D distribution of the photo-evoked activation. Postsynatpic potential is monitored in a cell of interest by whole-cell recording. (ii) Normalized response of two-photon-induced activation of CoChR (left y-axis, green) and ΔF/F0 of jRGECO1b (right y-axis, magenta) as a function of wavelength. Excitation wavelength for photostimulation (950 nm) was chosen based on the normalized curve. To minimize imaging-induced activation of opsin-expressing neurons and reach high Ca^2+^ sensitivity, imaging was done at 1140 nm. (iii) Optical setup for simultaneous volumetric imaging and single-cell stimulation. The stimulation path is shown in green. The imaging path is shown in magenta. The remote focusing unit is highlighted with a dotted rectangle. PBS, polarizing beam splitter; QWP, quarter wave plate; SLM, spatial light modulator; PMT, photomultiplier tube; M, mirror. **b.** A transgenic zebrafish expressing CoChR-EGFP-Kv2.1 (green) and jRGECO1b (magenta) in V2a neurons in the side view (top) and the dorsal view (bottom). d, dorsal; v, ventral; r, rostral; c, caudal. The dotted whited rectangle indicates the region show in the panel ei. **c.** (i) Experimental setup for assessing imaging-induced activation of opsin-expressing cells. Hindbrain V2a were co-labeled with CoChR-EGFP-Kv2.1 and jRGECO1b and their membrane potentials were monitored by perforated patch recordings. The patched cell was stimulated by two-photon holographic illumination. Ca^2+^ response was monitored by jRGECO1b imaging. An example trace of membrane potential (top, black trace), stimulation pulses (green dots, 2 mW, 20 ms, 5 Hz), and Ca^2+^ response (bottom, magenta). (ii) A close up view of the action potentials induced by stimulation pulses. (iii) Imaging-induced depolarization. The duration of the imaging is shown with the magenta rectangle. An arrow highlights the repolarization after the end of the imaging scan. (iv) Imaging-induced depolarization in the neurons that elicited action potentials in response to stimulation (n = 10 cells). **d.** Physiological measurement of spatial resolution of holographic stimulation in neurons expressing CoChR-EGFP-Kv2.1. (i) Whole-cell recordings were performed in hindbrain V2a neurons expressing CoChR-Kv2.1. Lateral resolution was examined by scanning the 6 μm diameter stimulation spot laterally by moving the sample in 4 μm step (left). A train of stimulation pulses (20 ms, 5 Hz) were delivered to induce spiking activity. The stimulation power was chosen to elicit reliable spiking across trials (4 mW). Axial resolution was similarly examined by moving the objective lens axially in 8 μm step (right). (ii) Normalized stimulation-induced spike counts as a function of distance from soma in lateral direction (orange, n = 14 cells) and in axial direction (cyan, n = 13 cells). (iii) Distribution of lateral and axial resolution (lateral, n = 14 cells; axial, n = 13 cells). Spatial resolution is defined as the full width at half maximum of the stimulation induced spike counts. **e.** Assessment of single-cell stimulation with volumetric jRGECO1b imaging. (i) The region of the hindbrain used for this experiment is indicated by a dotted white rectangle in the dorsal view (left) and the side view (top right). CoChR-EGFP-Kv2.1, green; jRGECO1b, magenta. Nine optical planes were acquired at 7-μm step at 6 Hz (bottom right, magenta cube). A 6-μm spot two-photon illumination was delivered in a 1-sec train of 20 ms pulses at 10 Hz (green). (ii) Examples of stimulations at the power just enough to activate the target cell (2 mW) and at the power that led to the activation of off-target cells (4 mW). Activated cells detected by regression analysis (see Methods) are shown in red based on their t-values on top of the dorsal view of the jRGECO1b volume. The activated cells and their jRGECO1b timecourses were shown on the right (thick magenta line, average response; thin magenta lines, individual responses). Z value indicates the offset in the axial direction in μm from the target cell. (iii) Spatial distribution of the targeted hindbrain V2a neurons (40 targets from 4 fish) and the activated cells at the condition that led to single-cell activation (36 targets). Red open circles indicate the targets with successful single-cell on-target activation (27 targets). Blue open circles indicate the targets with successful single-cell but off-target activation (9 targets). Black crosses indicate the targets with no successful single-cell activation (4 targets). The activated cells for successful single-cell activation are indicated by filled circles with their color coding the stimulation power. (iv) Stimulation power used for single-cell stimulation (n = 36 targets). (v) Number of activated cells as a function of the normalized power (normalized to the power used for single-cell stimulation for each target) (n = 36 targets). The data was fitted with a linear regression (black line). The regression coefficient was statistically significant (*P* = 2.6e-17). (vi) The location of activated cells relative to the target cell at the power for single-cell stimulation (cyan) and at the power just above the power for single-cell stimulation (orange) (n = 36 targets). The distribution changed significantly in the dorso-ventral axis (Kolmogorov-Smirnov test, *P* = 9.6e-3).

For our study, we chose the high-photocurrent channelrhodopsin CoChR^6^ based on previous successful demonstrations of two-photon cellular stimulation with this opsin^5,7^. To monitor activated neurons by Ca^2+^ imaging, we chose a genetic red Ca^2+^ indicator, jRGECO1b, which has been successfully used in zebrafish^8^. This indicator allowed the imaging of Ca^2+^ response at the wavelength that minimizes the imaging-induced activation of CoChR (Fig. 1a ii; Supplementary Fig. 1) and was sensitive enough to detect single action potentials in zebrafish sensory neurons (Supplementary Fig. 2). For two-photon activation of CoChR, we set up a two-photon patterned illumination system based on computer generated holography^5^ that allows the generation of photostimulation spots in arbitrary locations in 3D. We achieved axial resolution of 12 μm for a 6 μm-diameter spot of illumination (Supplementary Fig. 3), which is appropriate for single-cell stimulation of zebrafish neurons whose average diameter is 6.6 μm^9^. To perform concurrent volumetric jRGECO1b imaging, we implemented a remote-focusing system based on a remote movable mirror that is translated by a voice coil motor^10^. This technique enables us to change the imaging-plane without moving the objective lens, thus makes it possible to perform volumetric imaging without changing the axial position of photostimulation spots (Fig. 1 aiii). This system achieves axial resolution of less than 4.5 μm within the axial scanning range of 140 μm which is enough to obtain two-photon imaging at cellular resolution (Supplementary Fig. 4).

To reach cellular resolution for opsin stimulation, we fused CoChR with the soma-localization sequence from the mammalian potassium channel Kv2.1^3^ to target its expression to cell bodies. We then generated two UAS transgenic lines: one that expresses CoChR fused with the Kv2.1 sequence (hereafter referred as CoChR-EGFP-Kv2.1), and one that expresses jRGECO1b (Fig. 1b) to take advantage of stronger but sparser expression one can attain with the Gal4/UAS system. Using these optical setups and transgenic lines, we examined the amount of imaging-induced activation of CoChR in a practical experimental condition in hindbrain neurons (Fig. 1c). To monitor subthreshold activation of CoChR during the imaging of jRGECO1b expressed in cytosol, we used perforated patch recording so as to retain jRGECO1b in the cytosol while recording. For this analysis, we only included neurons with high CoChR expression that exhibited reliable action potentials in response to stimulation pulses (Fig. 1c i,ii) and used single-plane imaging with high enough imaging power (6.5 mW) to reliably detect the fluorescent increase of jRGECO1b (Fig. 1ci). As a given neuron is scanned more frequently with single-plane imaging, this test provides an upper estimate of the imaging artifact. We measured the hyperpolarization after the end of imaging to examine the imaging-induced activation at a steady state (Fig. 1ciii). The activation ranged from 0 to 2.1 mV with median of 0.4 mV (Fig. 1civ), showing that our optical configuration minimizes the chance of causing spurious off-target activations during single-cell stimulation or state changes of the imaged area.

**Supplementary Figure 1.**
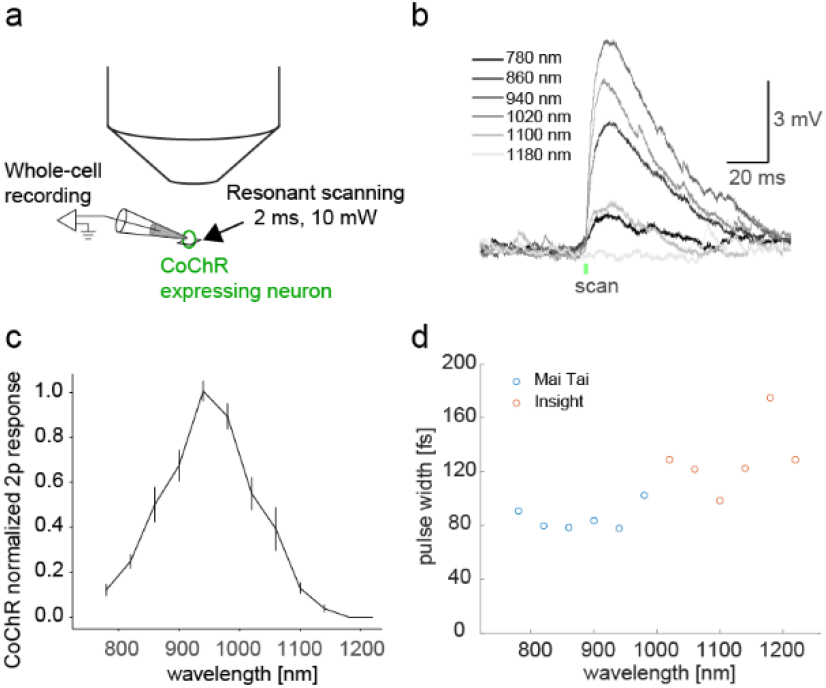
Wavelength-dependent two-photon activation of CoChR-expressing neurons. **a.** Experimental setup. Subthreshold response in a hindbrain V2a neuron expressing CoChR was monitored by whole-cell recording. The recorded cell was stimulated by resonant raster scanning of femtosecond laser over the cell body (2 ms, 10 mW). **b.** Example traces of subthreshold response from 780 nm to 1180 nm. The timing of the stimulation scanning is indicated by a green dot. **c.** Normalized response amplitude as a function of wavelength (n = 10 cells). **d.** Pulse widths of femtosecond laser as a function of wavelength. Measurement from two different laser sources are combined (blue, Mai Tai; red, Insight).

**Supplementary Figure 2.**
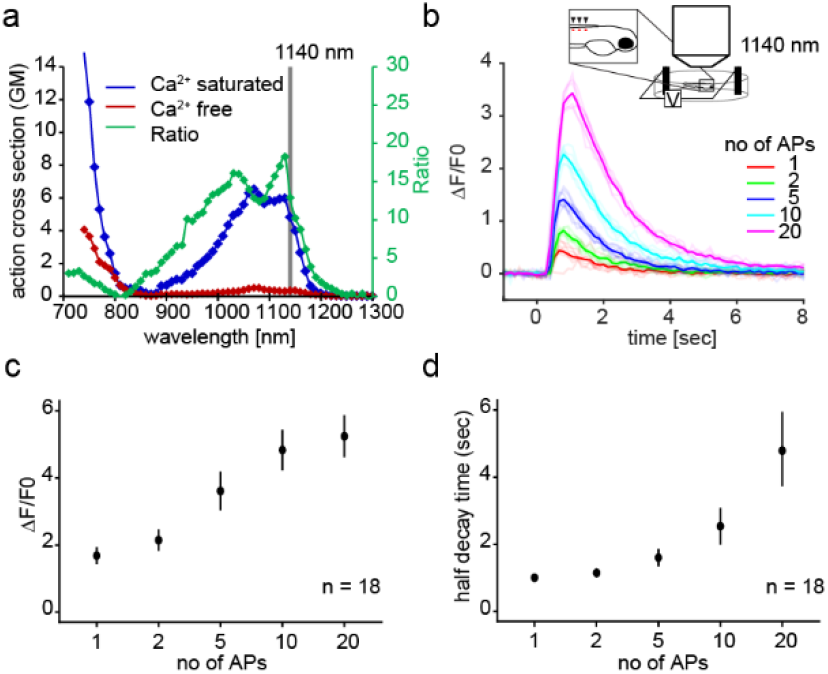
Sensitivity of jRGECO1b in zebrafish spinal sensory neurons. **a.** Two-photon action cross section of jRGECO1b in Ca^2+^-free (blue line) and Ca^2+^-saturated (red line) states. Green line shows the ratio between the two states. **b.** Two-photon imaging of zebrafish Rohon-Beard cells expressing jRGECO1b at 1140 nm. Example traces of jRGECO1b Ca^2+^ response to action potentials elicited by electrical shocks (50 Hz). **c.** Fluorescence fold change of jRGECO1b as a function of number of action potentials (APs). **d.** Half decay time of jRGECO1b as a function of number of action potentials (APs).

**Supplementary Figure 3.**
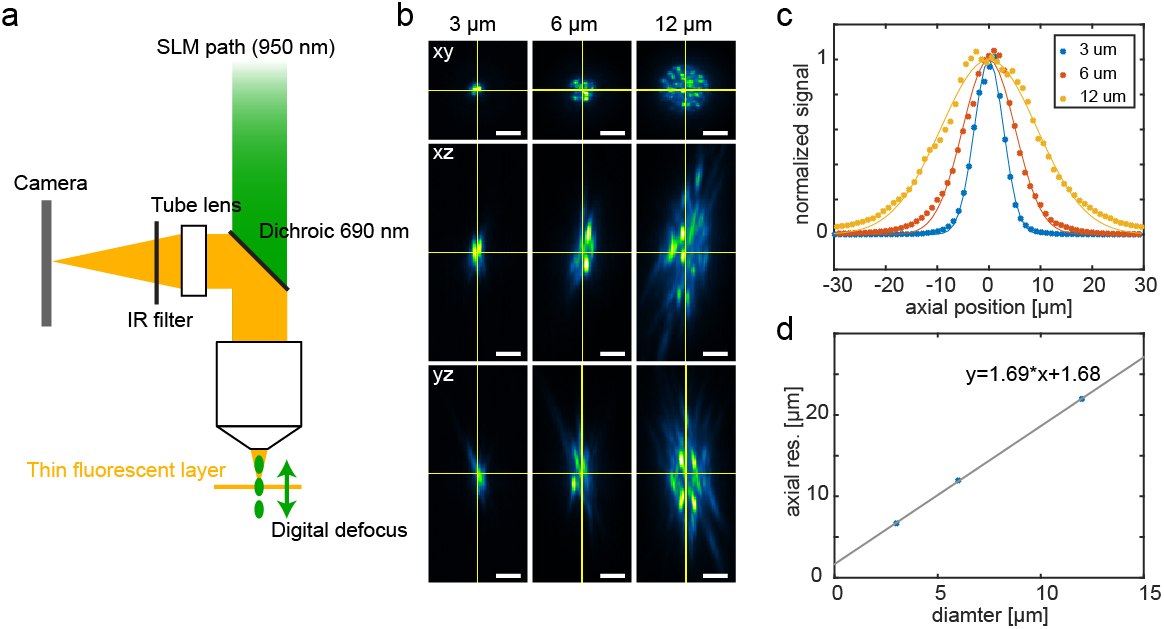
Axial profile of two-photon holographic stimulation. **a.** Experimental setup for characterizing axial resolution of two-photon holographic beam. A cover glass with a thin layer of fluorescent dye is brought to the focal plane of the objective lens. A holographic spot of a given diameter is digitally focused at different depths while imaging the resulting fluorescence at the focal plane of the objective lens with a camera. **b.** Cross sections of holographic spots with variable diameters (3, 6 and 12 μm). **c.** Axial profiles of the variably sized holographic spots (3, 6 and 12 μm). Normalized integrated fluorescent signal is plotted as a function of axial position. **d.** Axial resolution as a function of spot diameter, showing a linear relationship between them.

**Supplementary Figure 4.**
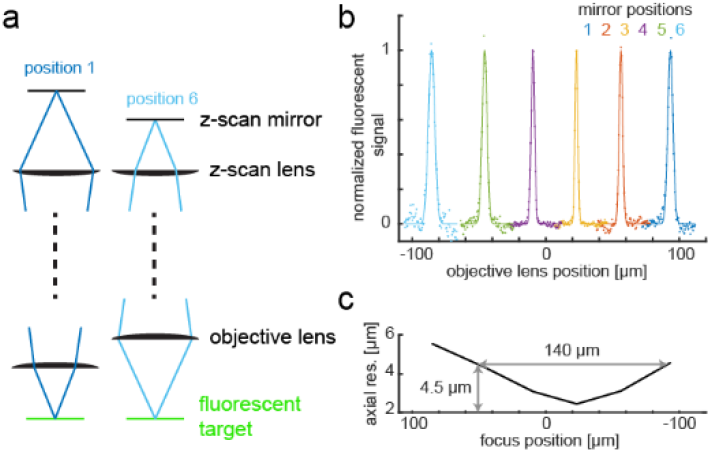
Axial resolution of the remote focusing system. **a.** Experimental procedure. Z-scan mirror was positioned by a voice coil to change the focus of the scanning beam (dark blue, position 1; cyan, position6). For each mirror position, we scan the objective lens axially and monitor the fluorescent signal from the fluorescent target (green) to characterize the axial extent of the point spread function (PSF). **b.** Normalized fluorescent signal as a function of objective lens position. Signal profiles were plotted separately for each mirror position (1 to 6). Zero in the x-axis (“objective lens position”) is defined as the position where the fluorescent target is in focus when the remote focusing unit is bypassed. **c.** Axial extent of the PSF as a function of the focus position. Axial extent is defined as full width at half maximum of the normalized fluorescent signal. Note that axial resolution stays below 4.5 μm for 140-μm axial scanning range.

Next, we examined the spatial resolution of our two-photon photoactivation system using two approaches. First, we used whole-cell recording to examine how spiking response to a 6-μm stimulation spot changes as a function of the offset between the stimulation spot and the target (Fig. 1d). The spatial resolution of stimulation based on spiking response was 6 μm in the lateral direction and 15 μm in the axial direction (n=12), which is on par with that of the optical confinement of the two-photon holographic illumination (Supplementary Fig. 3). We then examined if it was possible to achieve single-cell stimulation of hindbrain neurons using concurrent volumetric jRGECO1b imaging (Fig. 1e). We used a slice interval of 7 μm which is on par with the average soma size of neurons in larval zebrafish neurons^9^ so that we do not miss collateral activation of nearby neurons (Fig. 1e i). By detecting activated cells based on online regression analysis (see Methods), we tuned the stimulation power so that we stimulate only a single cell (Fig. 1e ii). Out of 40 targets we examined throughout the hindbrain population (Fig. 1e iii), we were able to find the power that activates just one cell in 36 cases, 9 of which activated an off-target cell presumably due to imperfect soma localization of CoChR by the Kv2.1 sequence^5^ and higher excitability and CoChR expression of the off-target cell. The power required for single-cell stimulation varied from 1.2 mW to 8 mW with median of 3.4 mW (Fig. 1e iv). The number of activated cells increased significantly when the power was set above the minimal power for single-cell stimulation (Fig. 1e v), further suggesting the importance of using the minimum power for each target for precise singe-cell and multi-cell activations. Specifically, collateral activations observed at suprathreshold conditions showed substantial spread in the axial direction (Fig. 1e vi; Kolmogorov-Smirnov test, *P* = 0.0096). This is probably due to the fact that higher excitation powers, approaching the saturation of the opsin response, axially enlarge the effective volume of the optical stimulation, as previously discussed^5^. This further suggests the importance of concurrent volumetric imaging with cellular z resolution to assess collateral activation (Fig. 1e i). Taken together, we showed that it is possible to attain single-cell stimulation when the stimulation power was carefully tuned using the combination of two-photon opsin stimulation, sparse opsin expression, and crosstalk-free volumetric functional imaging.

To demonstrate the utility of this system for mapping synaptic connectivity, we focused on the investigation of the connectivity from spinal V2a neurons to axial primary motoneurons (PMNs). Spinal V2a neurons are glutamatergic ipsilateral projecting neurons that play a critical role in locomotion^11^. Their connectivity to motoneurons has been examined with both paired whole-cell recordings and electron microscopy^12–14^. However, these analyses have been restricted to local connections within a few spinal segments. To examine their long-range connectivity, we patched PMNs in spinal segment 23 and stimulated single V2a neurons expressing CoChR-Kv2.1 and jRGECO1b from segment 6 to segment 21 (Fig. 2a i, n = 6 fish). In 85% of the stimulation sites, we were able to achieve single-cell activation (358/419; Fig. 2a ii). Twenty seven percent of the cells we activated elicited membrane depolarization in the recorded PMNs in synchrony with the stimulation pulses (98/358; Fig. 2b i, ii). We found two distinct response types based on the kinetics of membrane depolarization that are not mutually exclusive. The first type had a fast rise time constant and was observed in response to 12% of the stimulated cells (Fig. 2b i; half rise time = 0.57 ± 0.35 ms, n = 43). This time constant is compatible with those of postsynaptic potentials^14^, indicating they are synaptically connected to the recorded PMNs. The second type had a slower rise time constant and was observed in response to 21% of the stimulated cells (Fig. 2b ii; half rise time = 15.3 ± 5.4 ms, n = 74). As motoneurons do not express CoChR in this preparation, the second type is likely to be indirect excitation through electrical coupling among axial motoneurons^14–16^.

**Figure 2.**
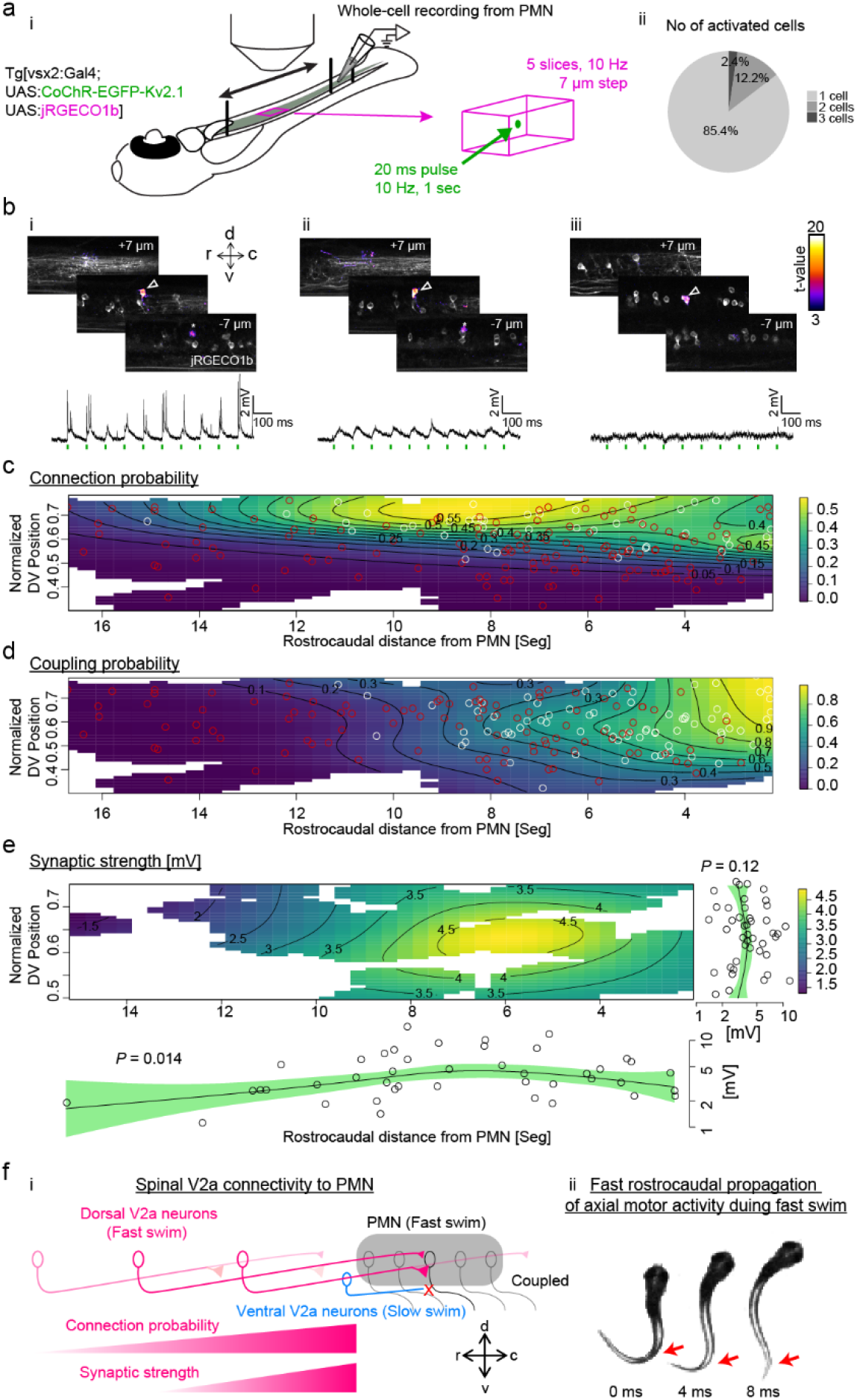
Large-scale synaptic connectivity mapping from spinal V2a neurons to primary motoneurons assisted by two-photon optogenetics and imaging. **a.** (i) Experimental procedure. Whole-cell recording was established in a primary motoneuron (PMN) in the caudal spinal cord (spinal segment 23) in the transgenic line expressing CoChR-EGFP-Kv2.1 and jRGECO1b in V2a neurons (indicated by the shaded area). Spinal V2a neurons rostral to the patched PMN were activated one-by-one by holographic two-photon stimulation. Simultaneous volumetric imaging of jRGECO1b was performed to examine the resulting activation. To cover V2a neurons up to 17 segments rostral to the patched PMN, the sample was scanned along the rostrocaudal direction. (ii) Number of activated V2a neurons per photostimulation sites (n = 419 sites, N = 6 fish) **b.** Examples of successful single-cell stimulation of spinal V2a neurons. (i) An example of V2a neuron activation that led to fast membrane depolarizations in a PMN. Three sagittal slices showing the activated cells (open arrowhead) and nearby neurons and neuropil. The activated pixels were detected by regression analysis and color-coded by their t-values. The target cell is indicated by an open arrow. The asterisk at the plane −7 μm indicates the part of the target cell. d, dorsal; v, ventral; r, rostral; c, caudal. The membrane potential from the patched PMN during the stimulation is plotted at the bottom. The individual stimulation pulses are indicated by green bars. Note the fast depolarizations triggered by stimulation pulses, indicating the stimulated V2a neuron is synaptically connected to the PMN. (ii) An example of V2a neuron activation that led to slow membrane depolarization in the PMN. This panel is similarly organized to i. Note that the stimulation pulses triggered slow depolarizations distinct from synaptic potential. (iii) An example of V2a neuron activation that led to no discernible responses in the PMN. No consistent postsynaptic potentials were observed in sync with the stimulation pulses. **c.** Spatial probability map of V2a neurons synaptically connected to PMN. The positions of the singly stimulated V2a neurons are plotted. The x-axis indicates the distance from the patched PMN in the caudal spinal cord in the unit of segments and the y-axis indicates the dorsoventral position normalized to the full thickness of the cord at a given segment (see Methods). Synaptically connected cells are indicated by white circles and non-connected ones are indicated by red circles. The probability density was estimated by generalized additive model (see Methods) and color coded. Both dorsoventral and rostrocaudal positions showed significant effect to the connection probability (dorsoventral position, *P* = 6.5e-6; rostrocaudal position, *P* = 0.015). **d.** Spatial probability map of V2a neurons exhibiting putative indirect electrical coupling (slow depolarization) to PMN. This panel is similarly organized to c. White circles, coupled cells; red circles, non-coupled cells. The rostral caudal position showed significant effect (*P* = 7.5e-9). **e.** Spatial distribution of the synaptic strength from the synaptically connected V2a neurons to PMN. This panel is similarly organized to c and d except that the partial effect of the dorsoventral position is plotted on the right and that of the rostrocaudal position is plotted on the bottom. Note the significant effect of rostrocaudal position. **f.** A proposed circuit motif of dorsal spinal V2a neurons involved in fast locomotion that entails fast rostrocaudal propagation of axial muscle activity. (i) PMNs are only recruited during fast swim and electrically coupled across local spinal segments (gray shade). PMNs do not receive inputs from ventral V2a neurons that are recruited during slow swim (blue) but do receive long-range inputs from dorsal V2a neurons that are recruited during fast swim (magenta). Both the connection probability and the synaptic strength decline as the distance between the dorsal V2a neurons and PMNs increases. This creates a long-range gradient of excitatory drive in the rostrocaudal direction, which is suitable for generating fast rostrocadual propagation of axial muscle activity. (ii) Fast rostrocaudal propagation of axial muscle activity during fast swim. Consecutive images of fish during fast swim are shown with the time interval of 4 ms. The regions of the tail where the curvature changes its sign are indicated with red arrows in all images to highlight the rostrocaudal propagation of axial motor activity during fast swim.

Coexistence of both response types were observed in 18% of the responding cell (18/98). The probability of finding synaptically connected cells was significantly biased towards the dorsal spinal cord (Fig. 2c; *P* = 6.5e-6). The bias in the rostrocaudal direction was also significant but less prominent with some connected cells up to 15 segments rostral to the PMN (Fig. 2c; *P* = 0.015). This not only confirms previous observations about the selective innervation of dorsal V2a neurons to PMNs^12,13^ but also extends this connectivity pattern to long-range connections. The probability of finding coupled cells was significantly biased to the nearby segments with more than half of the cells within 6 segments from the recorded PMNs (Fig. 2d; *P* = 7.5e-9) in agreement with a previous report^14^. The distribution of synaptic strength also showed significant bias in the rostrocaudal direction with stronger connections from the closer V2a neurons (Fig. 2e; *P* = 0.014). Such a rostrocaudal gradient of excitatory drive has been suggested to play an important role in the rostrocaudal propagation of axial muscle activity^17,18^ and was also hypothesized to arise from either more abundant excitatory neurons in the rostral spinal cord^18,19^ or their more prominent descending projections^17,20^. Here we provide experimental evidence for long-range rostrocaudal gradients of synaptic strength and connection probability from a class of excitatory interneurons that is selectively recruited during fast locomotion^21,12^ (Fig. 2f i). This evidence suggests the role of this circuit motif in fast undulatory movements during rapid forward locomotion. (Fig. 2f ii).

In summary, we established a system that enables the efficient discovery of cellular-level circuit motifs *in vivo*. The combination of circuit optogenetics and concurrent volumetric Ca^2+^ imaging assures single-cell precision optical activation of putative presynaptic neurons. High temporal resolution of electrical recording of postsynaptic response enables the identification of different types of postsynaptic response based on its kinetics. Thus, this system allows single-cell precision optical mapping of presynaptic neurons and detailed characterization of synaptic dynamics on a large scale and enables efficient discovery of circuit motifs as we demonstrated here. Temporally precise multi-spot photoactivation also enables the investigation of synaptic summation. This approach, combined with rapidly improving sensitive and fast voltage indicators, can pave the way for an all-optical synaptic mapping to assess longitudinal changes in synaptic properties across many temporal scales in a living animal.

## Methods

### Fish care

Zebrafish larvae were obtained from an in-house breeding colony of adults maintained at 28.5°C on a 14-10 hr light-dark cycle. Embryos were raised in a separate incubator but at the same temperature and on the same light-dark cycle. Larvae at 3 and 5 days post-fertilization (dpf) were used for this study. All experiments presented in this study were conducted in accordance with the animal research guidelines from the National Institutes of Health and were approved by the Institutional Animal Care and Use Committee and Institutional Biosafety Committee of Janelia Research Campus (16-145).

### Transgenic fish

The following previously published transgenic lines were used in this study: TgBAC(vsx2:Gal4)^22^; Tg(10xUAS:CoChR-EGFP)^*jf44* 8^; Tg(elavl3:Gal4)^23^. Tg(10xUAS:CoChR-EGFP-Kv2.1) and Tg(10xUAS:jRGECO1b) were generated with the Tol2 system (Urasaki, Morvan, and Kawakami 2006) using the corresponding plasmids prepared from the following reagents.

1. Tol2kit^24^
2. FCK-gene86-GFP (CoChR)^6^
3. pTol2-elavl3-Voltron-ST(Kv2.1)^25^
4. pTol2-elavl3-jRGECO1b^8^

### Optics

Two-photon computer generated holography and remote-focusing systems were built on top of a custom microscope equipped with a two-photon resonant scanning system and a one-photon widefield epifluorescence system as described previously^25,26^. Briefly, a resonant scanner (Thorlabs, MPM-2PKIT) was used to scan the femtosecond laser (MKS Instruments, Mai Tai HP Deep See and Insight Deep See, MA) through a scan lens assembly consisting of three achromatic lenses (Thorlabs, AC254-150-C, AC508-200-B and AC508-200-B), a tube lens (Nikon, MXA22018) and a 25x objective lens (Leica, HCX IRAPO L 25x/0.95 W, Germany). Excited fluorescence by two-photon scanning was directed to GaAsP photomultiplier tubes (Hamamatsu, H10770-40 SEL) by a dichroic mirror (Semrock, FF735-Di02) and spectrally separated to three channels by dichroic mirrors (Semrock, FF562-Di03 and FF650-Di01) and emission filters (Semrock, FF01-520/70-30-D, FF01-625/90, and FF01-731/137). Two-photon images were collected using ScanImage (Vidrio Technologies, VA). Epi-fluorescence for widefield imaging was directed to the path by an image-splitting dichroic mirror (Semrock, F640-FDi02-t3) and filtered by an IR filter (Semrock, FF01-750sp) and an appropriate emission filter, and then images were formed on the camera (PCO, pco.edge 4.2, Germany) through a tube lens (Nikon, MXA22018). Images were collected using μManager (https://micro-manager.org/). The initial dichroic mirror for each path is on a miniature stage to switch between the two detection pathways.

### Computer generated holography system

Two-photon illumination patterns were obtained with computer-generated holography, a technique based on the phase modulation of the laser wavefront by the use of a spatial light modulator (SLM) ^5^. The beam from a femtosecond laser (MKS instruments, Mai Tai HP, Deep See, MA) was enlarged by a telescope and reflected off the SLM (Hamamatsu, LCOS-SLM X10468-07) and then projected on the back focal plane of the objective lens with a double afocal telescope (f= 30 mm, Thorlabs AC254-030-B; f= 100 mm, Thorlabs AC508-100-B; f = 200 mm, Thorlabs AC508-200-B; f = 300 mm, Thorlabs AC508-300-B). The zeroth order of diffraction was eliminated by introducing a single cylindrical lens (Thorlabs, LJ1516L1)^27^. The photoactivation beam were directed to the objective lens through a dichroic mirror (Semrock, Di02-R1064). A phase profile for a given illumination pattern was calculated using the Gerchberg and Saxton (GS) iterative algorithm^28,29^, implemented on a custom-designed software (Wave Front Designer). Lens-phase modulations were added to 2D-phase holograms to enable remote axial displacement and 3D positioning of 2D light patterns. The performance of the illumination system for single cell photostimulation was assessed by the reconstruction of the illumination volumes corresponding to circular holographic patterns (diameter: 3 μm, 6 μm and 12 μm) (Supplementary Fig. 3). A thin layer of rhodamine-6G (spin coated and dried on a glass coverslip) was positioned at the focal plane of the objective lens and then a disk illumination pattern was scanned axially by the lens-phase modulation and the resulting fluorescence was imaged by the widefield epifluorescence imaging setup described above (Supplementary Fig. 3a). The fluorescence signal was integrated over the spot surface and plotted as a function of the axial position specified by the lens-phase modulation (Supplementary Fig. 3c). The effective axial shift induced by the nominal digital phase-modulation was verified in a dual microscope configuration by a mechanical axial scan of the objective lens, as previously described^27,5^.

The holographic illumination system was calibrated before each experiment by verifying the spatial alignment between the patterns of illumination and the imaging system. We generated a bi-dimensional multi-spot holographic pattern (typically 5 diffraction limited spots) and we used it to bleach a thick fluorescence plate (Chroma, part No. 92001). The pattern was axially shifted by a lens-phase modulation in order to bleach the fluorescent plate at different depths (typically five planes over ~200 μm along the z axis). Then we re-imaged the bleached pattern with the two-photon scanning imaging system to obtain the exact positioning of the holographic spots in the coordinates of the imaging system. An affine transformation (x-y-z stretch, translation and rotation) between the nominal and the measured position of the spots was identified for each plane. The parameters of such transformation were linearly interpolated to obtain the required transformation matrix at any depth.

After such calibrations, we examined the targeting accuracy by bleaching the fluorescent plate with an arbitrary multi-point 3D illumination patterns (Supplementary Fig. 5), ensuring micrometric precision in the control of the spot locations.

**Supplementary Figure 5.**
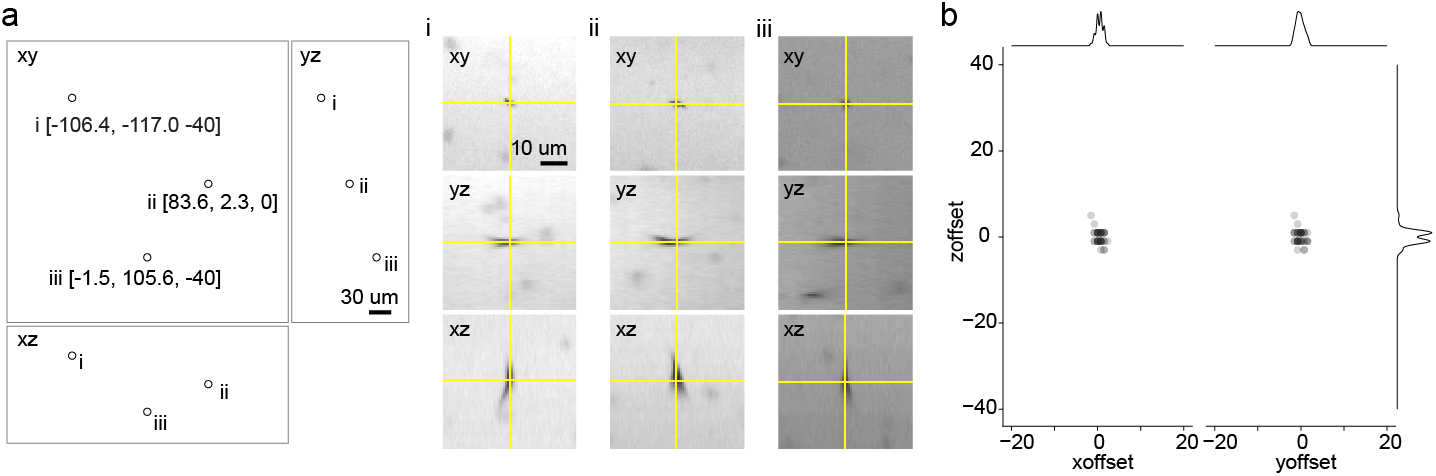
Accuracy of 3D targeting in vitro tested by bleaching fluorescent plate. **a.** Examples of bleached spots. (left) Location of three bleached spots in xy, yz, and xz views. (right) Close-up views of bleached spots. **b.** The location of bleached spots relative to the target location (n = 42 targets).

### Remote-focusing system

An axial scanning system for two-photon imaging was implemented based on a remote movable mirror that is conjugate to the specimen plane and translated by a voice coil motor similar to the system described previously^10^ (Figure 1a iii). This system is built on top of the existing two-photon imaging system in a way that can be engaged on demand using the polarizing beam splitters (Thorlabs, PBS203) mounted on a sliding stage. When the beam is directed to the system, it goes through a telescope (f1 = 75 mm Thorlabs AC254-075-C, f2 = 125 mm Thorlabs AC254-125-C) and an axial scan lens assembly consisting of three achromats lenses (Thorlabs AC254-100-C, AC254-75-C and AC254-100-C) before being reflected by the axial scan mirror (Thorlabs, PF03-03-P01) mounted on a voice coil (equipment solutions, LFA-2004). A quarter-wave plate (Thorlabs, AQWP05M-980) was inserted in the path close to the polarizing beam splitter so that the beam reflected by the mirror goes past the polarizing beam splitter and to the xy-scanning module. Basic optical performance was assessed by measuring the point spread function (Supplementary Fig. 4). We chose 6 mirror positions and in each position we took a z-stack of a two-photon excitable target by moving the objective lens by a piezo objective scanner (PI, P-725K129, Germany) with a 0.5 μm step size (Supplementary Fig. 4a). The signal profile was plotted as a function of the position of the objective lens for each mirror position (Supplementary Fig. 4b). Then the full width at half maximum for each mirror position was plotted as a function of the focus position (Supplementary Fig. 4c).

### Assessment of the imaging-induced artifact by simultaneous perforated patch recording and Ca^2+^ imaging from holographically stimulated neurons

Perforated patch recordings were made in hindbrain neurons expressing CoChR-EGFP-Kv2.1 and jRGECO1b in Tg[vsx2:Gal4;10xUAS:CoChR-EGFP-Kv2.1;10xUAS:jRGECO1b] using the procedure described previously with some modifications^26^. Briefly, 5 dpf fish were paralyzed by 1 mg/ml alpha-bungarotoxin (MilliporeSigma, 203980, MO) dissolved in system water and then anesthetized with MS-222 and secured with etched tungsten pins through the notochord to a glass-bottom dish coated with Sylgard (Dow Corning, Sylgard 184, MA). Then the solution was exchanged with extracellular solution containing MS-222 (134 mM NaCl, 2.9 mM KCl, 1.2 mM MgCl2, 2.1 mM CaCl2, 10 mM HEPES, 10 mM glucose, 0.3 mM MS-222, adjusted to pH 7.8 with NaOH). Then the ventral surface of the hindbrain was carefully exposed using a tungsten pin and fine forceps and the solution was exchanged to extracellular solution without MS-222. Ventral neurons in the middle hindbrain (hindbrain segment 3, 4 and 5) were targeted based on fluorescence and scanned Dodt gradient contrast image acquired with the custom two-photon microscope. Nystatin-based perforated whole-cell recordings were established using the procedure described previously. Briefly, nystatin-containing intracellular solution was freshly prepared from 50 μL intracellular solution (125 mM K-gluconate, 2.5 mM MgCl2, 10 mM EGTA, 10 mM HEPES and 4 mM Na2ATP adjusted to pH 7.3 with KOH) and 250 nL nystatin stock solution (100 mg/mL in DMSO) and filtered by 0.45 μm nanopore filter (Ultra free MC 0.45μm). Then a glass pipette with the resistance of 7-10 MΩ was front-filled with 2 μL intracellular solution and then backfilled with 14 μL nystatin-containing intracellular solution. After quickly forming a GΩ seal with the target cell, we waited until the access resistance became lower than 100 MΩ. Then, the membrane potential was recorded in current-clamp mode with EPC10 Quadro amplifier (HEKA Electronik, Germany) using PatchMaster software (HEKA Electronik, Germany) while simultaneously delivering a train of holographic stimulation (6 μm circular pattern, 950 nm, 10 pulses of 20-ms duration delivered at 40 Hz) and recording Ca^2+^ response with two-photon jRGECO1b imaging (1140 nm, 6.5 mW, 30 Hz). Imaging power was set to ensure Ca^2+^ response was reliably detected. Only the cells that showed reliable firing in response to the holographic stimulation were included in the analysis of imaging-induced depolarization. The amount of depolarization by the imaging laser was quantified by calculating the difference in the average membrane potential from the three-second window before the end of the imaging and that from three-second window after the end of the imaging (10 trials).

### Characterization of wavelength-dependent two-photon activation of CoChR

Current-clamp recording was performed from hindbrain neurons expressing CoChR-EGFP-Kv2.1 in Tg[vsx2:Gal4; 10xUAS:CoChR-EGFP-Kv2.1] using the procedure described above but in a standard whole-cell configuration. The patched cell was stimulated by scanning femtosecond laser with resonant-galvo mirrors (6 μm x 6 μm, 2 ms duration) using two femtosecond laser sources to cover from 780 nm to 1220 nm in 40 nm steps (780-980: MaiTai HP; 1020-1220: Insight HP). The power after the objective lens was matched to 10 mW across wavelengths so that the none of the cells spiked an action potential across all the wavelengths examined. The pulse width at each wavelength was measured with auto-correlator (FEMTOCHROME Research, Inc., FR-103TPM/700). The resulting membrane depolarization to the most effective wavelength for a given cell was averaged from at least 5 trials and then used to normalize the response amplitude in individual trials. This normalized response was averaged over all the trials pooled across cells. The resulting action spectrum is similar to published recordings^5^, that here we extend to a broader wavelength range.

### Characterization of jRGECO1b in zebrafish sensory neurons

The sensitivity of jRGECO1b in zebrafish sensory neurons was examined as described previously^30^. Briefly, Tg[elavl3:Gal4;10xUAS:jRGECO1b] at 3 dpf were paralyzed by a 5-min bath application of 1 mg/ml a-bungarotoxin (Sigma, 203980). Larvae were mounted on their side in a field stimulation chamber (Warner, RC-27NE2) with 1.5 % low-melting-point agarose (Supplementary Fig. 2b). Tactile sensory neurons (Rohon-Beard cells) were identified based on their oblong-shaped soma in the dorsal spinal cord and their innervations to the skin using the two-photon microscope described above. Functional images (256 × 256 pixels) were acquired at 7.5 Hz (1140 nm, 3 mW) while trains of 1, 2, 5, 10, and 20 field stimuli (1 ms pulse width at 50 Hz) were applied with a stimulator (NPI, ISO-STIM) to stimulate the processes of the sensory neurons on the skin. This protocol elicits the corresponding number of spikes in the sensory neurons^30^. The stimulation voltage was calibrated to elicit an identifiable response to a single pulse in Rohon-Beard cells without stimulating muscle cells. Regions of interest were selected manually, and data were analyzed using MATLAB (MathWorks). Two-photon absorption spectra of jRGECO1b were collected by John Macklin as previously described^31^ (Supplementary Fig. 2a).

### Concurrent two-photon optogenetics and volumetric imaging

Tg[vsx2:Gal4;10xUAS:CoChR-EGFP-Kv2.1;10xUAS:jRGECO1b] at 5 dpf were used for this experiment. The fish with high expression of CoChR-EGFP-Kv2.1 and jRGEOC1b were selected, paralyzed with α-bungarotoxin (MilliporeSigma, 203980, MO), and then embedded in 1.6% low-melting point agar. Each sample was left under the objective lens at least 40 minutes before starting the experiment to minimize potential sample drifts during the experiment. A reference z-stack for jRGECO1b was acquired (1140 nm, 512 x 512 x 100 slices, 0.76 x 0.76 x 2 μm). Then a single target cell was selected for each stimulation session. A phase pattern for stimulating each target with a 6-μm diameter disk pattern was calculated, taking into account the transformation matrix obtained in the calibration described above. A train of ten 20 ms stimulation pulses was delivered at 10 Hz while monitoring Ca^2+^ response by volumetric imaging of jRGEOC1b using the remote-focusing system (1140 nm, 512×256, 9 planes, 7 μm z step, 6 Hz, 6.5 mW) from 5 s before the stimulation to 14 s after the stimulation. To identify neurons stimulated by this protocol during the experiments, we performed regression analysis as described previously^26^. Briefly, two-photon data was first corrected for mismatch between the odd and even scan lines by cross correlation and then the x-y movements were corrected for each slice by cross correlation if necessary. A regressor for stimulation-induced activity was constructed by convolving the stimulation pulses with a jRGECO1b impulse response function modeled as the rise and decay exponentials (0.5 s rise and 2 s decay) and then standardized. The standardized coefficient was estimated by ordinary least square and then *T* value for each pixel was calculated based on standardized coefficient and residual. The activation map was used to determine the soma position of all the activated cells and their distances to the intended targets were calculated using the transform from the remote-focusing imaging space to the standard imaging space as described above. We repeated this procedure with stimulation powers ranging from 1.2 mW to 12 mW to find the condition where only one cell is activated for each stimulation target. ΔF/F0 was calculated by defining the baseline fluorescence (F0) as the average fluorescence during the 4-s time window before the stimulation.

### Synaptic connectivity mapping of spinal V2a neurons to spinal motoneurons

Whole-cell recordings from spinal primary motoneurons were performed from Tg[vsx2:Gal4;10xUAS:CoChR-EGFP-Kv2.1;10xUAS:jRGECO1b] at 5 dpf using the procedure described previously^26^. Briefly, the fish with high expression of CoChR-EGFP-Kv2.1 and jRGEOC1b were selected, paralyzed with α-bungarotoxin, and pinned to a sylgard-coated dish with etched tungsten pins. The spinal cord at muscle segment 23 was exposed by removing the muscles overlying the spinal cord. A pipette was filled with the standard intracellular solution containing 0.01% of Alxea Fluor 647 hydrazide (Thermo Fisher Scientific) and a class of primary motoneurons, caudal primary motoneuron (CaP), was targeted based on its large soma and proximity to the segmental boundary and the lateral surface of the spinal cord. The stimulations of putative presynaptic spinal V2a neurons started 10 minutes after forming a gigaohm seal so that the fluorescence of Alexa Fluor 647 does not leak to the red channel used for jRGECO1b imaging. Single spinal V2a neurons were targeted by a 6 μm-diameter disk illumination pattern. Ten 20-ms stimulation pulses were delivered at a 100-ms interval so that postsynaptic potentials induced by each pulse can be separated temporally. Ca^2+^ responses of neighboring V2a neurons were monitored by jRGECO1b imaging with the remote focusing system (512×256, 5 planes, 7 μm step, 10 Hz). The stimulation session was continued until the patched CaP neuron started to depolarize more than 5 mV after establishing the whole-cell configuration. Then a series of z-stacks for CoChR-EGFP-Kv2.1, jRGECO1b and Alexa Fluor 647 was acquired in the standard two-photon imaging configuration from muscle segments 4 to 23.

The regression analysis was performed to detect all the cells activated by the stimulation using the procedure mentioned above but with one modification. To isolate the stimulation induced Ca^2+^ response from the Ca^2+^ response during spontaneous swimming, an additional regressor for spontaneous swimming was constructed based on the slow subthreshold depolarization of PMNs observed during spontaneous swimming^21^. Such depolarization was detected by a bandpass filter and then the regressor was constructed by convolving a boxcar function representing the duration of swim with the jRGECO1b impulse response function described above. The trials with more than one activated cell were excluded from further analysis. Among the stimulation trials with only one activated cell, we detect postsynaptic potentials that occurred within 20 ms from the onset of each stimulation pulse based on the spiking pattern we observed in hindbrain neurons using the identical stimulation protocol. If such postsynaptic potentials occurred in more than 5 stimulation pulses for a given activated cell and their rise times matched the rise times of synaptic potentials^14^, the cell was considered to have synaptic connection to the patched motoneuron (half rise time = 0.57 ± 0.35 ms, n = 43). In some cases, slow depolarization with a much longer rise time was observed consistently after the stimulation pulses (half rise time = 15.3 ± 5.4 ms, n = 74). Such slow membrane depolarization eliciting input to primary motoneurons was previously characterized and suggested to be caused by electrical coupling through neighboring motoneurons^14^. Thus, the cells that produced such slow response in motoneurons were categorized as electrically coupled cells. To reconstruct the position of each activated cell, a series of z-stacks was stitched with the stitching plugin available through Fiji (https://fiji.sc/).The activated cell for each stimulation trial was identified in this stitched z-stack based on the x-y position of the stage and the z position of the objective lens for each trial, and further confirmed by manual inspection. In order to pool data from multiple fish, the positions of the activated cells were transformed to a common space using a procedure similar to the one described previously^21^. First, for a given rostrocaudal position, the midline was identified based on the presence of central canal in Dodt gradient contrast image, and the dorsal edge and ventral edge of the spinal cord were marked using segmented lines in Fiji. This procedure was repeated to define the dorsal and ventral edges of the spinal cord from muscle segments 4 to 23. Then the distances from a given activated cell to the nearest line segments for the dorsal and the ventral edge were calculated. Then the dorsoventral position of a given activated cell was defined as *Distance_ventral_*/(*Distance_dorsal_* + *Distance_ventral_*). To define the rostro-caudal position within each segment, muscle segment boundaries were marked with single-segment lines in Fiji. Then a line was extended from a given activated cell to the nearest segmental boundaries so that the vector of this line is the vector sum of the nearest dorsal and ventral edges. Then the normalized rostro-caudal position of a given activated cell within a segment was defined as *Distance_caudal_*/(*Distance_rostral_* + *Distance_caudal_*). The rostral caudal position of a given activated cell relative to the patched CaP was defined as the sum of this normalized rostro-caudal position within a segment and number of segments from the caudal end of the muscle segment where the recorded CaP was located to the caudal end of the segment where a given activated cell was located. Then the two dimensional distributions of synaptic connection probability, coupling probability and synaptic strength were modeled with generalized additive models using the *mgcv* package (https://cran.r-project.org/web/packages/mgcv/) in R (https://www.r-project.org/). This statistical technique is widely used to model spatial data in ecology and epidemiology^32^. Probability distributions were modeled as a combination of a full tensor product smooth (*te*) of the normalized dorsoventral position, a full tensor product smooth (*te*) of the rostrocaudal position, and a tensor product interaction (*ti*) of the normalized dorsoventral position and the rostrocaudal position with a logit link function for binomial distribution. Synaptic strength was modeled similarly but with log transformed to satisfy the assumptions of the linear model (normality, homogeneity of variance, independence and linearity). The tensor product interaction was discarded to improve the concurvity of the model^32^ (the worst case for the full model: 0.99; the worst case for the model without the tensor product interaction: 0.38).

## Acknowledgements

We thank A. Pujala and M. Tanimoto for critical feedback and helpful comments on the manuscript; J. Macklin and GENIE project for providing two-photon absorption spectra of jRGECO1b; V. de Sars for the development of the Wave Front Designer software for holographic phasemask calculation; V. Goncharov for help with our optical system. This work was supported by Howard Hughes Medical Institute, the National Institute of Health (Grant NIH 1UF1NS107574 - 01), the Fondation Bettencourt Schueller (Prix Coups d’élan pour la recherché française) and the Getty Lab.

## Author contributions

C.M. designed and built the remote-focusing system. D.T. designed and built the holographic stimulation system. L.Z. generated Tg[10xUAS:CoChR-EGFP-Kv2.1], and managed the transgenic lines used in the study. C.Y. and M.B.A. generated Tg[10xUAS:jRGECO1b]. M.K., C.M., and D.T. designed and performed experiments, and interpreted data. M.K., D.T., and V.E. wrote the paper. M.K. and V.E. conceived and supervised the project.

